# Subpopulations of stressed *Y. pseudotuberculosis* preferentially survive doxycycline treatment within host tissues

**DOI:** 10.1101/2020.04.13.039222

**Authors:** Jasmine Ramirez Raneses, Alysha L. Ellison, Bessie Liu, Kimberly M. Davis

## Abstract

Severe systemic bacterial infections result in colonization of deep tissues, which can be very difficult to eliminate with antibiotics. It remains unclear if this is because antibiotics are not reaching inhibitory concentrations within tissues, if subsets of bacteria are less susceptible to antibiotics, or if both contribute to limited treatment efficacy. To determine the concentration of doxycycline (Dox) present within deep tissues following treatment, we generated a fluorescent transcriptional reporter derived from the *tet* operon to specifically detect intracellular tetracycline exposure at the single bacterial cell level. Dox exposure was detected in the spleen 2 hours after intraperitoneal injection, and by 4 hours post-injection, this treatment resulted in a significant decrease in viable *Yersinia pseudotuberculosis* in the spleen. Nitric oxide-stressed bacteria preferentially survived treatment, suggesting stress was sufficient to alter Dox susceptibility. Many bacteria (~10%) survived a single dose of Dox, and the antibiotic accumulated at the periphery of microcolonies to growth inhibitory concentrations until 48 hours post-treatment. After this timepoint, antibiotic concentrations decreased and bacterial growth resumed. Dox-treated mice eventually succumbed to the infection, albeit with significantly prolonged survival relative to untreated mice. These results indicate that Dox delivery by intraperitoneal injection results in rapid diffusion of inhibitory concentrations of antibiotic into the spleen, but stressed cells preferentially survive drug treatment, and bacterial growth resumes once drug concentrations decrease. This fluorescent reporter strategy for antibiotic detection could easily be modified to detect the concentration of additional antimicrobial compounds within host tissues following drug administration.

**Importance:** Bacterial infections are very difficult to treat when bacteria spread into the bloodstream and begin to replicate within deep tissues, such as the spleen. Subsets of bacteria can survive antibiotic treatment, but it remains unclear if this survival is because of limited drug diffusion into tissues, or if something has changed within the bacteria, promoting survival of some bacterial cells. Here, we have developed a fluorescent reporter to detect doxycycline (Dox) diffusion into host tissues, and show that Dox impacts the bacterial population within hours of administration, and inhibits bacterial growth for 48 hours. However, bacterial growth resumes when antibiotic concentrations decrease. Subsets of bacteria, stressed by the host response to infection, survive Dox treatment at a higher rate. These results provide critical information about the dynamics that occur within deep tissues following antibiotic administration, and suggests subsets of bacteria are predisposed to survive inhibitory concentrations of antibiotic before exposure.

## Introduction

Bacterial infections are exceedingly difficult to treat with antibiotics when bacteria spread systemically into deep tissues. It is thought that this difficulty arises because the antibiotic cannot accumulate to inhibitory concentrations within these tissue sites, so bacteria are exposed to sub-inhibitory concentrations of the drug. It has been difficult to determine the antibiotic concentration present within deep tissues, and many studies have relied on serum concentrations to determine whether tissue contain inhibitory levels (1, 2). Recently, mass spectrometry-based assays have shown differential diffusion of tuberculosis drugs into granulomas, where rifampicin and pyrazinamide can penetrate past the edge of granulomas, but moxifloxacin does not diffuse into the caseum (3). These results suggest select drugs could penetrate deep tissues efficiently, but further studies are needed to determine the diffusion of additional drug compounds in the context of other infections. Mass spectrometry requires specialized technology for tissue processing, data collection, and analysis, which can be cost prohibitive. Bacterial fluorescent reporters offer a cost-effective alternative to detect antibiotic concentrations within tissues and individual bacterial cells, and provide spatiotemporal information about antibiotic concentrations following treatment.

Slow growing bacterial cells are known to be less susceptible to antibiotics, but it remains unclear which pathways promote slowed bacterial growth within host tissues (4, 5). Nutrient limitation may reduce metabolic activity, and subsequently reduce antibiotic susceptibility (6–8). Host immune cell subsets can promote the formation of slow growing bacteria within host tissues (9, 10), but it remains unclear if antimicrobial compounds, such as reactive oxygen species (ROS) or reactive nitrogen species (RNS), contribute to slowed growth (11, 12). Recently, it was shown that ROS exposure was sufficient to reduce the rifampicin susceptibility of *S. aureus* during spleen infection (13). However, it remains unclear if bacterial cells responding to ROS preferentially survived drug treatment. Expression of virulence factors, specifically type-III secretion system (T3SS) expression, has been linked to slowed growth and decreased antibiotic susceptibility in *Salmonella*, but this link has not been made within host tissues (14, 15).

*Yersinia pseudotuberculosis* replicates within deep tissue sites to form clonal extracellular clusters of bacteria called microcolonies (or pyogranulomas) (16–18). Within the spleen, neutrophils are recruited to sites of bacterial replication and directly contact bacteria at the periphery of microcolonies. Peripheral bacteria respond to neutrophil contact by expressing high levels of the type-III secretion system (T3SS), which promotes translocation of T3SS effector proteins into neutrophils and inhibits phagocytosis. A layer of monocytes is recruited immediately around the layer of neutrophils. Recruited monocytes develop characteristics of dendritic cells and macrophages (16–18), and expresses inducible nitric oxide synthase (iNOS), which produces antimicrobial nitric oxide (NO) that diffuses towards the microcolony. Bacteria at the periphery of the microcolony respond to NO by expressing the NO-detoxifying gene, *hmp* (18). *hmp* expression at the periphery prevents NO diffusion into the center of microcolonies, establishing an example of cooperative behavior, where the peripheral bacteria protect the interior bacteria from the antimicrobial action of NO (18, 19). It remains unclear if the protective expression of *hmp* by peripheral cells comes at a fitness cost, and if this stress response is sufficient to alter the antibiotic susceptibility of this subpopulation.

The genetic tools available for *Y. pseudotuberculosis* allow us to easily construct fluorescent reporters to detect changes in the host environment at the single bacterium level (18–20). Here, we introduce a novel tetracycline-responsive fluorescent reporter based on the *tet* operon, that enables detection of the diffusion of tetracycline derivatives, including doxycycline, within host tissues. Doxycycline was chosen for these studies because it is a relevant bacteriostatic antibiotic for treatment of *Yersinia* infection (21–23), but similar reporters could also be generated for other classes of antibiotics. This tool allows us to determine if individual bacteria are differentially exposed to antibiotics during drug treatment within mouse tissues and more closely approximate antibiotic concentrations within host tissues.

## Results

### Characterizing the doxycycline-responsive reporter

Bacterial infections of deep tissues are exceedingly difficult to treat, and it is thought this is because the concentration of antibiotic does not reach inhibitory levels within deep tissue sites. To detect antibiotic diffusion and exposure within bacterial microcolonies, we constructed a tetracycline-responsive transcriptional reporter within a low copy plasmid (pMMB67EH), and transformed the reporter into wild-type (WT) *Y. pseudotuberculosis*. The reporter contains a constitutively expressed *tetR* gene, whose gene product, TetR, binds to DNA and represses expression of *P*_*tetA*_::*mCherry* in the absence of tetracycline derivatives. In the presence of tetracyclines, drug binding to TetR relieves repression, and transcription of *mCherry* occurs (Figure 1A).

**Figure 1:**
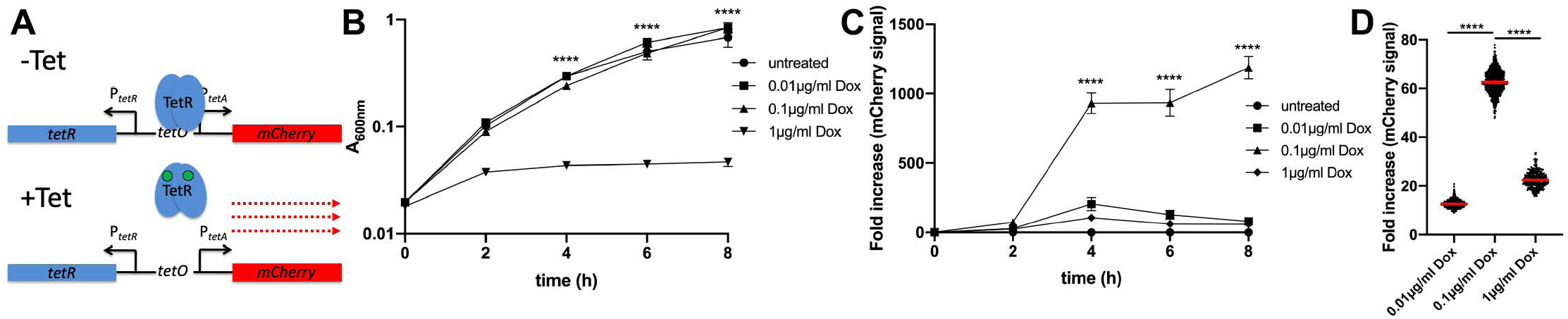
Characterizing the doxycycline-responsive reporter. A) Tetracycline responsive reporter schematic. In the absence of tetracyclines (-Tet), the repressor (TetR) binds at the *tetO* sequence to inhibit expression of *mCherry*, inserted downstream of the *P*_*tetA*_ promoter. In the presence of tetracyclines (+Tet), TetR repression is relieved, and *mCherry* is expressed. B) Growth curve of the *P*_*tetA*_::*mCherry* strain with the indicated doses of Dox. Optical density (A_600nm_) is measured over time (hours, h). Mean and error of three biological replicates are shown. C) Reporter expression during growth curve, detected with 560nm excitation/610nm emission, expressed as fold increase mCherry signal relative to untreated cells. Median and range depicted, four biological replicates. D) Reporter expression within individual bacterial cells detected by fluorescence microscopy after 4h treatment, expressed as fold increase in mean mCherry signal/cell relative to average signal of untreated cells. Experiment performed in triplicate, one representative is shown with medians. Statistics: B) & C) Two way ANOVA, Tukey’s multiple comparisons test, D) Kruskal-Wallis with Dunn’s multiple comparison test, ****p<0.0001.

To determine if bacteria containing the *P*_*tetA*_ reporter can be used to quantify the amount of doxycycline (Dox) exposure within individual bacteria, we treated liquid cultures of the reporter strain with increasing doses of Dox. Bacterial growth was quantified by absorbance (A_600nm_) and mCherry fluorescence was quantified in a microplate reader. Treatment with 1µg/ml Dox resulted in significant growth inhibition relative to untreated cells, 0.01µg/ml Dox, and 0.1µg/ml Dox (Figure 1B), which is consistent with previously published Dox minimal inhibitory concentrations (MIC) levels for *Yersinia* (24–26). Reporter signal was detected with 0.01µg/ml Dox, and significantly increased with 0.1µg/ml Dox; neither lower dose inhibited bacterial growth (Figure 1B, 1C). 1µg/ml exposure resulted in decreased reporter expression, likely due to ribosomal inhibition by Dox. Reporter signal was first detected after 2 hours (h) of antibiotic exposure and significantly increased after 4h of exposure (Figure 1C).

Plate reader measurements give averaged, population-level readings, and cannot detect heterogeneity across individual cells. To determine if bacterial responses to Dox were heterogeneous or uniform, individual bacterial cells were also imaged by fluorescence microscopy to quantify reporter signal at the single-cell level. Single cell quantification of reporter signal was consistent with the population-level results, where 0.01µg/ml treatment increased reporter signal, 0.1µg/ml resulted in a significant increase in signal, and 1µg/ml treatment decreased reporter signal within individual cells (Figure 1D). Cells within each treatment group had distinct levels of fluorescent signal in response to Dox, suggesting this reporter will be a useful tool for determining antibiotic exposure levels. Collectively, these results confirm the reporter responds to Dox, and can be used to detect antibiotic exposure within individual bacterial cells.

### Doxycycline differentially diffuses across microcolonies

To determine if Dox can diffuse into microcolonies and impact bacterial growth, we infected C57BL/6 mice intravenously with GFP^+^ *P*_*tetA*_::*mCherry Y. pseudotuberculosis*, and injected mice intraperitoneally with 40mg/kg Dox at 48h p.i. The 48h timepoint p.i. was chosen to allow microcolonies to initially replicate and form distinct bacterial subpopulations prior to antibiotic treatment (18); intraperitoneal administration of 40mg/kg Dox should result in inhibitory levels of Dox in the serum (1-4µg/ml) within hours of injection (27, 28). Spleens were harvested at 24h post-treatment to quantify CFUs, microcolony areas, and reporter expression (Figure 2A). Dox treatment resulted in significantly fewer CFUs relative to untreated mice, however ~10^5^ CFUs remained in the spleen 24h after treatment (Figure 2B). The area of microcolonies was also significantly lower following 24h of Dox treatment compared to untreated mice, suggesting antibiotic treatment either inhibited growth of microcolonies, or promoted elimination of a subset of bacteria (Figure 2C).

**Figure 2:**
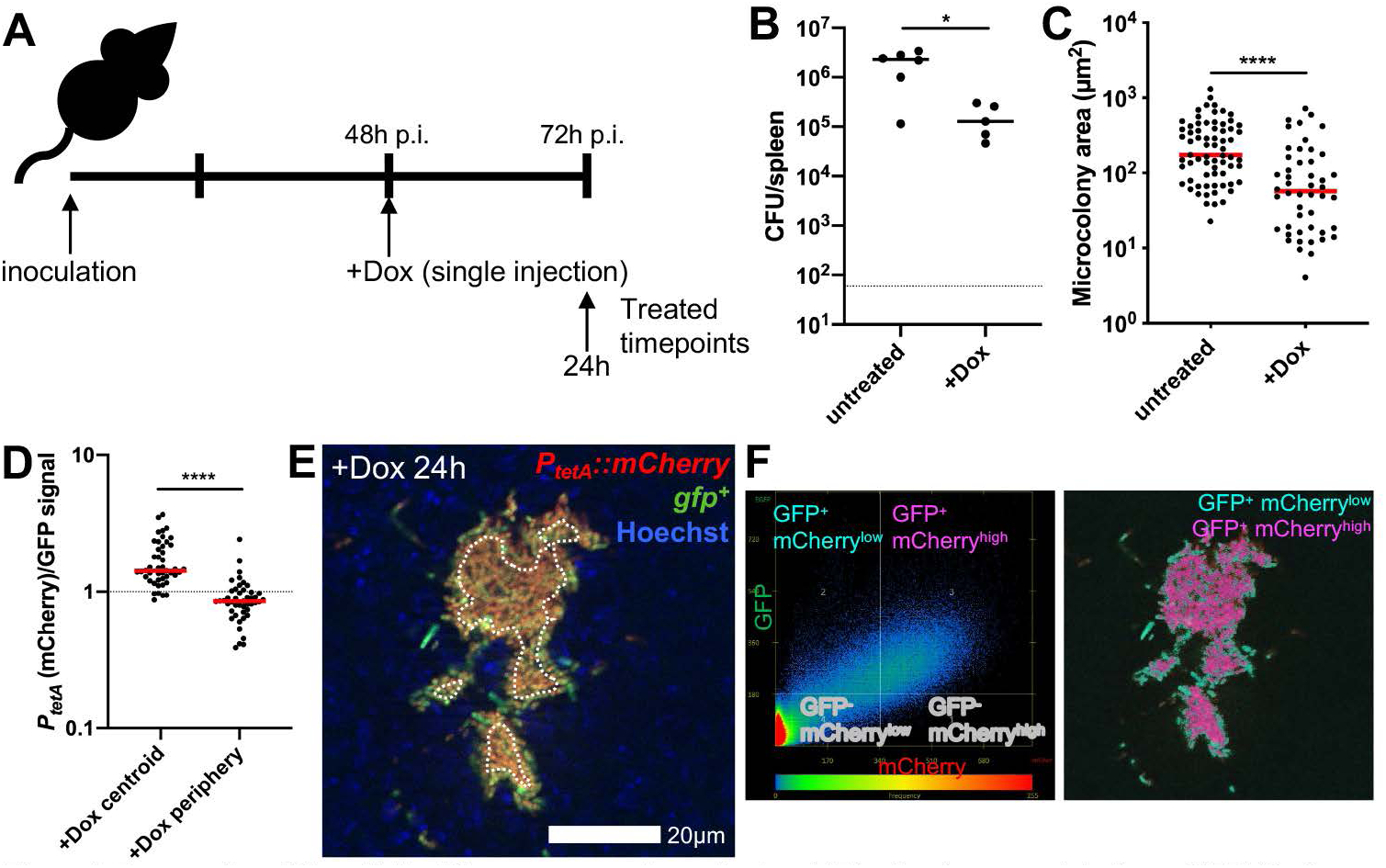
Doxycycline differentially diffuses across microcolonies. A) Timeline for mouse infections. C57BL/6 mice were inoculated intravenously (i.v.) with 10^3^ GFP^+^ *P*_*tetA*_::*mCherry Y. pseudotuberculosis*. Mice were treated at 48h post-inoculation (p.i.) with 40mg/kg Dox (Dox) in 100 µl PBS, administered intraperitoneally (i.p.), and spleens were harvested at the indicated timepoints to quantify CFUs or process tissue for fluorescence microscopy. B) CFU/spleen quantification from treated (+Dox, 24h) or untreated mice (72h). Dots: individual mice. C) Fluorescence microscopy of spleens to quantify microcolony area (µm^2^) from treated (+Dox) or untreated mice, based on GFP signal. Dots: individual microcolonies. D) Reporter signals were quantified from peripheral and centroid cells within treated microcolonies in the spleen, reporter signal was divided by GFP. E) Representative image, cells outside dotted white line are defined as peripheral. F) Co-localization analysis of image in E, to show differential reporter signal by pseudocolor. Dot plot depicts signal intensity for each pixel within the image. Quadrants were set using GFP^+^ signal to distinguish bacteria from surrounding tissue, within this gate, the 66% highest mCherry pixels are shown as mCherry^high^ (fuschia), and lowest 34% mCherry pixels are shown as mCherry^low^ (teal). Mouse infections were completed in three distinct experiments, animal numbers depicted in Figure 2B were processed for fluorescence microscopy in 2C-2F, medians are shown. Statistics: B) & C) Mann-Whitney, D) Wilcoxon matched-pairs, *p<0.05, ****p<0.0001

To determine if Dox diffused into microcolonies, the Dox-responsive reporter signal was compared to a constitutively-expressed GFP. Reporter signal was significantly higher at the centroid of microcolonies compared to the periphery (outside dotted white line), suggesting Dox diffused into the microcolony, and that there was differential diffusion of Dox across microcolonies at 24h post-treatment (Figure 2D, 2E). Co-localization analysis was used to highlight differences in mCherry signal intensity across microcolonies, and shows high mCherry reporter expression in the middle of microcolonies (fuschia) and low mCherry signal within peripheral cells (teal, Figure 2F). Decreased reporter signal at the periphery may indicate that heightened levels of antibiotic accumulate at this location, limiting the translational activity of peripheral cells. However, it could also suggest antibiotic accumulation at the centroid of microcolonies and lower concentrations at the periphery.

### Tetracycline derivatives accumulate at the periphery of microcolonies

Our initial experiments in Figure 2 utilized a reporter constructed with a stable mCherry fluorescent protein, which means reporter signal could accumulate over the course of the experiment. To confirm reporter signal represents current responses to Dox, we generated a fluorescent reporter construct with a destabilized mCherry fluorescent protein. The destabilized mCherry within this construct has a *ssrA* tag, which targets mCherry for proteasomal degradation and quickens protein turnover (29, 30). The *ssrA*-tagged mCherry has a half-life of 35.5 minutes (Figure 3A), which will allow detection of recent responses to the antibiotic. Liquid cultures of the destabilized reporter strain were treated with increasing doses of Dox, and fluorescent reporter expression was quantified at 4h post-treatment in a plate reader. Reporter expression patterns for the destabilized reporter were similar to the stable reporter at 4h; destabilized fluorescent signal was maximal with 0.1µg/ml Dox (Figure 3B).

**Figure 3:**
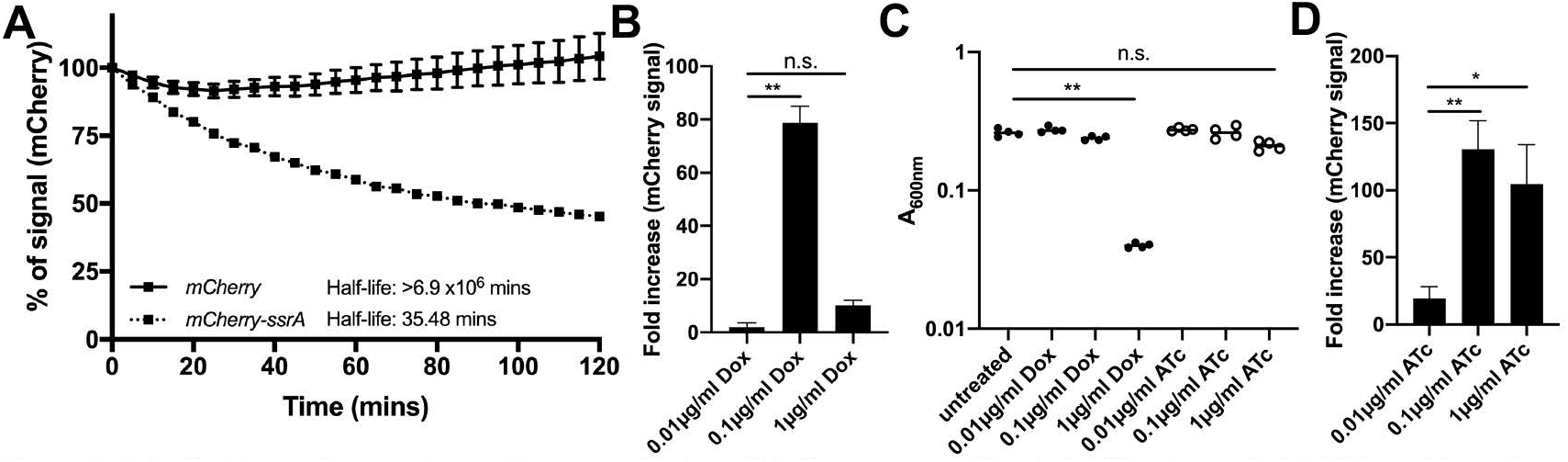
Anhydrotetracycline can be used to approximate antibiotic exposure without significant growth inhibition. A) One-phase exponential decay equations calculate the half-life of mCherry and destabilized mCherry-ssrA. Fluorescence expression was induced with IPTG, and kanamycin was then added to inhibit further translation. Fluorescence was detected over time (minutes). Data represent % of fluorescent signal intensity relative to time 0 (kanamycin addition). Mean and standard deviation of 6 biological replicates shown. B) Unstable fluorescent reporter (*P*_*tetA*_::*mCherry-ssrA*) expression with the indicated doses of Dox, treated for 4h at 37° C, fluorescence detected by plate reader. Fold increase relative to untreated cells, 4 biological replicates shown with median and range. C) Optical density (A_600nm_) at 4h post-treatment with the indicated doses of Dox and ATc. Dots represent individual cultures. D) Unstable fluorescent reporter expression with the indicated doses of ATc, treated for 4h at 37° C, fluorescent detected by plate reader. Fold increase relative to untreated cells, 5 biological replicates shown with median and range. Statistics: Kruskal-Wallis with uncorrected Dunn’s, *p<0.05, **p<0.01. n.s.: not significant.

To determine if the Dox concentrations at the periphery of microcolonies are reaching levels that inhibit translation, we performed experiments with the inactive tetracycline derivative, anhydrotetracycline. Anhydrotetracycline (ATc) can bind TetR and de-repress the *tet* operon, however, relative to Dox, ATc has ~35x lower binding affinity for *E. coli* ribosomes (31, 32). As it remained unclear if high concentrations of ATc impact *Y. pseudotuberculosis* growth, we treated liquid cultures of the reporter strain with increasing doses of ATc and detected growth and fluorescence with a microplate reader. Growth was significantly reduced with 1µg/ml Dox but was not significantly impacted by 1µg/ml ATc (Figure 3C). Reporter signal was detected with 0.01µg/ml ATc, and significantly increased with 0.1µg/ml ATc. However, 1µg/ml ATc also resulted in high levels of reporter expression, in contrast to results with Dox, suggesting ATc does not significantly inhibit ribosomal activity at this dose (Figure 3D). Similar patterns were seen using flow cytometry to detect fluorescent reporter signals (Supplemental Figure 1). The high level of reporter expression with 1µg/ml ATc suggested we could use ATc treatment to determine if peripheral bacteria are exposed to higher levels of antibiotic than the centroid, as increasing doses of ATc results in heightened reporter signal.

We then used the destabilized reporter strain in combination with Dox or ATc treatment to determine the concentration of antibiotics within microcolonies, to determine when antibiotic exposure occurs after treatment, and to confirm spatial differences in reporter signal are not due to accumulation of stable fluorescent protein. Mice were infected intravenously with the GFP^+^ *P*_*tetA*_::*mCherry-ssrA* strain, treated at 48h p.i., and splenic tissue was harvested at 2h, 4h, and 24h post-treatment (Figure 4A). Reporter expression was significantly higher at the centroid of microcolonies after 24h of Dox treatment, suggesting differential reporter expression is not due to accumulation of stable fluorescent protein (Figure 4B). Reporter signal was detected 2h after Dox injection, similar to exogenous addition in culture, indicating bacteria are exposed to the antibiotic soon after intraperitoneal injection (Figure 4B). During ATc treatment, reporter expression was significantly higher at the periphery of microcolony compared to the centroid (Figure 4C). Based on *in vitro* reporter expression (Figure 3B, 3D), these results suggest peripheral bacteria are exposed to inhibitory levels of ~1µg/ml tetracyclines, while the interior is exposed to lower sub-inhibitory concentrations, around 0.1µg/ml.

**Figure 4:**
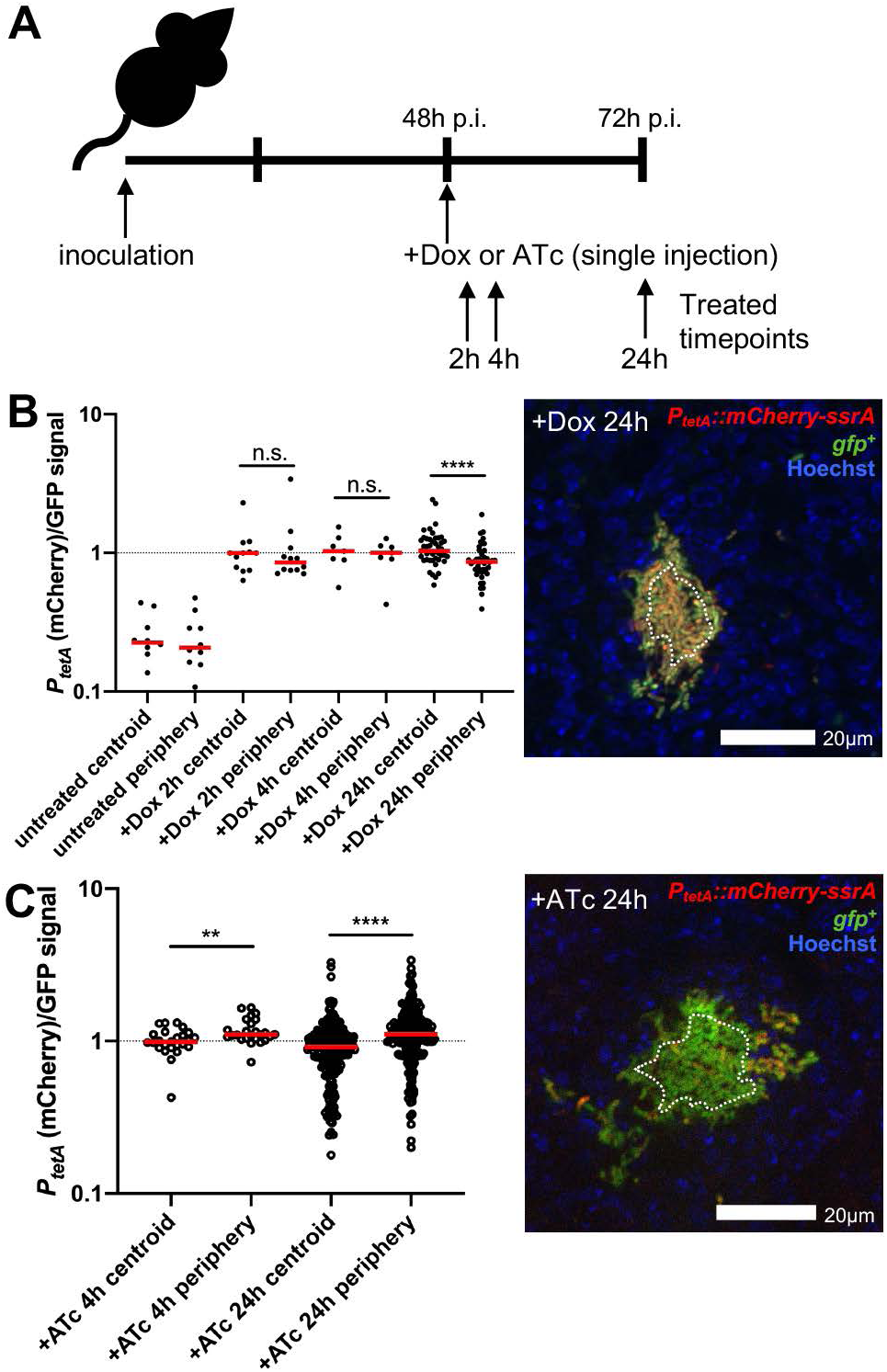
Tetracycline derivatives accumulate at the periphery of microcolonies. A) Timeline for mouse infections. C57BL/6 mice were inoculated i.v. with 10^3^ GFP^+^ *P*_*tetA*_::*mCherry-ssrA Y. pseudotuberculosis*. Mice were treated at 48h p.i. with 40mg/kg Dox or 40mg/kg ATc or untreated (48h), spleens were harvested at the indicated timepoints post-treatment to process for fluorescence microscopy. B) Reporter signal was quantified relative to GFP within peripheral and centroid cells, within microcolonies following Dox treatment. Dotted line: value of 1 (equivalent fluorescent intensity). Dots: individual microcolonies. Experiments performed on two days (4-5 mice/group), a representative image is shown (cells outside dotted white line are considered periphery). C) Reporter signal was quantified within peripheral and centroid cells within microcolonies following ATc treatment. Experiments performed on two distinct days (5-6 mice/group), a representative image is shown (cells outside dotted white line are considered periphery). Medians shown. Statistics: B) & C) Wilcoxon matched-pairs, **p<0.01, ****p<0.0001. n.s.: not significant.

### Microcolonies significantly decrease in size after doxycycline treatment

Dox was first detected within host tissues at 2h post-treatment (Figure 4B), but it remained unclear how antibiotic administration impacted bacterial growth and viability. To address this question, the splenic tissue depicted in Figure 4 was also harvested to quantify CFUs and microcolony areas. CFUs decreased 2h after Dox treatment, and there was a significant reduction in CFUs at 4h of Dox treatment relative to ATc (Figure 5A). The difference in CFUs was more pronounced at 24h post-treatment, where Dox treatment resulted in significantly fewer CFUs compared to ATc treatment or untreated mice, and ATc treatment CFUs were similar to untreated (72h) mice (Figure 5A). This suggested the difference in CFUs was due to antibiotic activity. There was a significant reduction in microcolony areas between 4h and 24h of Dox treatment, while microcolonies areas increased slightly during ATc treatment, suggesting Dox activity led to the decrease in microcolony areas (Figure 5B). Consistent with this, at 24h post-treatment, Dox treatment resulted in significantly smaller microcolony areas than ATc treatment or untreated (72h) mice (Figure 5B). ATc treatment also resulted in slightly smaller microcolonies than untreated tissue, suggesting there may have been some impact of ATc administration. These results suggest that many bacteria are eliminated following a single Dox treatment, while subsets of viable bacteria also remain within tissues at 24h after treatment.

**Figure 5:**
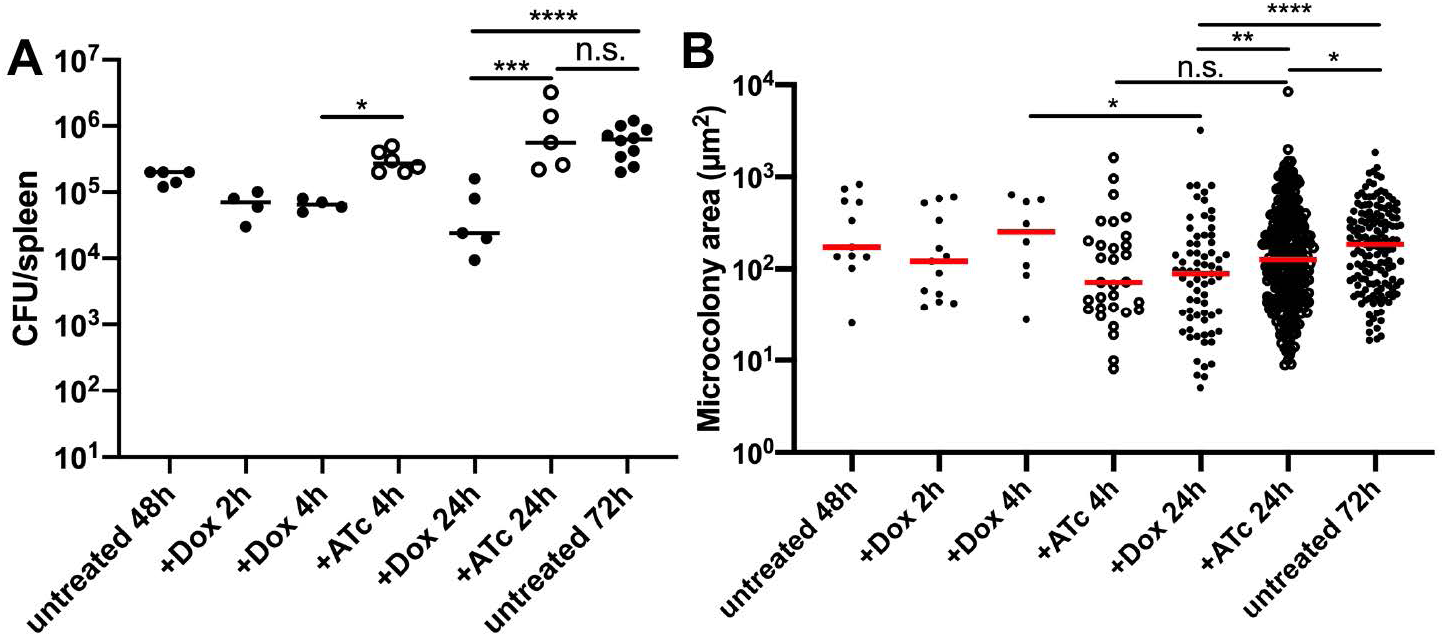
Microcolonies significantly decrease in size after doxycycline treatment. C57BL/6 mice were inoculated i.v. with 10^3^ GFP^+^ *P*_*tetA*_::*mCherry-ssrA Y. pseudotuberculosis*. Spleens were harvested at 48h p.i., additional mice were treated at 48h p.i. with Dox or ATc, and spleens were harvested at the indicated timepoints post-treatment to quantify CFUs and microcolony areas. A) CFU/spleen quantification from treated or untreated mice. Dots: individual mice. Experiments performed in triplicate. B) Microcolony area (µm^2^) quantification by fluorescence microscopy, from treated or untreated mice, based on GFP signal. Dots: individual microcolonies. Medians shown. Statistics: Kruskal-Wallis one-way ANOVA with uncorrected Dunn’s test, *p<0.05, **p<0.01, ***p<0.001, ****p<0.0001. n.s.: not significant.

### Hmp^+^ cells preferentially survive doxycycline treatment

We hypothesized that peripheral bacterial cells stressed by the host immune response may have reduced metabolic activity, and may be predominant within the surviving bacterial population after Dox treatment. The 48h treatment timepoint was specifically chosen throughout this manuscript to allow the stressed bacterial subpopulation to develop prior to treatment, to then determine if stress impacts survival of this subpopulation. To determine whether Hmp^+^ cells preferentially survive Dox treatment, we infected C57BL/6 mice intravenously with GFP^+^ *hmp::mCherry Y. pseudotuberculosis*, treated mice with Dox or ATc at 48h p.i., and splenic tissue was harvested at 2h and 4h post-treatment to quantify CFUs and quantify the proportion of Hmp^+^ bacteria by flow cytometry. CFUs in the spleen were significantly lower with 2h and 4h of Dox treatment compared to ATc treatment, suggesting antibiotic activity contributed to decreased CFUs (Figure 6A). Hmp^+^ bacteria were detected by flow cytometry, by gating on the total GFP^+^ bacterial population in the spleen, and quantifying the % Hmp^+^ cells before and after treatment. Mice were infected with the GFP^+^ strain in parallel to define the threshold for mCherry^+^ cells. The % Hmp^+^ cells significantly increased after 2h of Dox treatment compared to 2h ATc treatment, suggesting that more of the Hmp^+^, stressed cells were surviving Dox treatment than the Hmp-, unstressed cells, and that this was due to the activity of Dox (Figure 6B). At 4h post-treatment, the % Hmp^+^ cells increased significantly with Dox treatment relative to ATc treatment and untreated mice, again suggesting Hmp^+^ cells were preferentially surviving drug treatment. Increased *hmp* expression was not a response to Dox treatment; Dox treatment did not significantly impact *hmp* transcript levels *in vitro* (Figure 6C) or in the mouse spleen (Figure 6D). Dox treatment caused a trend towards decreased *hmp* transcript levels both *in vitro* and *in vivo*, although statistically this was not significant. We also confirmed that exposure to high levels of Dox (1µg/ml) does not increase the mCherry signal of *Y. pseudotuberculosis* by flow cytometry (Supplemental Figure 1), again suggesting increased *hmp* reporter signal is due to preferential survival of Hmp^+^ cells. Collectively, these results indicate that the stressed Hmp^+^ bacteria at the periphery of microcolonies are less susceptible to Dox than the Hmp^−^ nonstressed bacteria at the interior of microcolonies.

**Figure 6:**
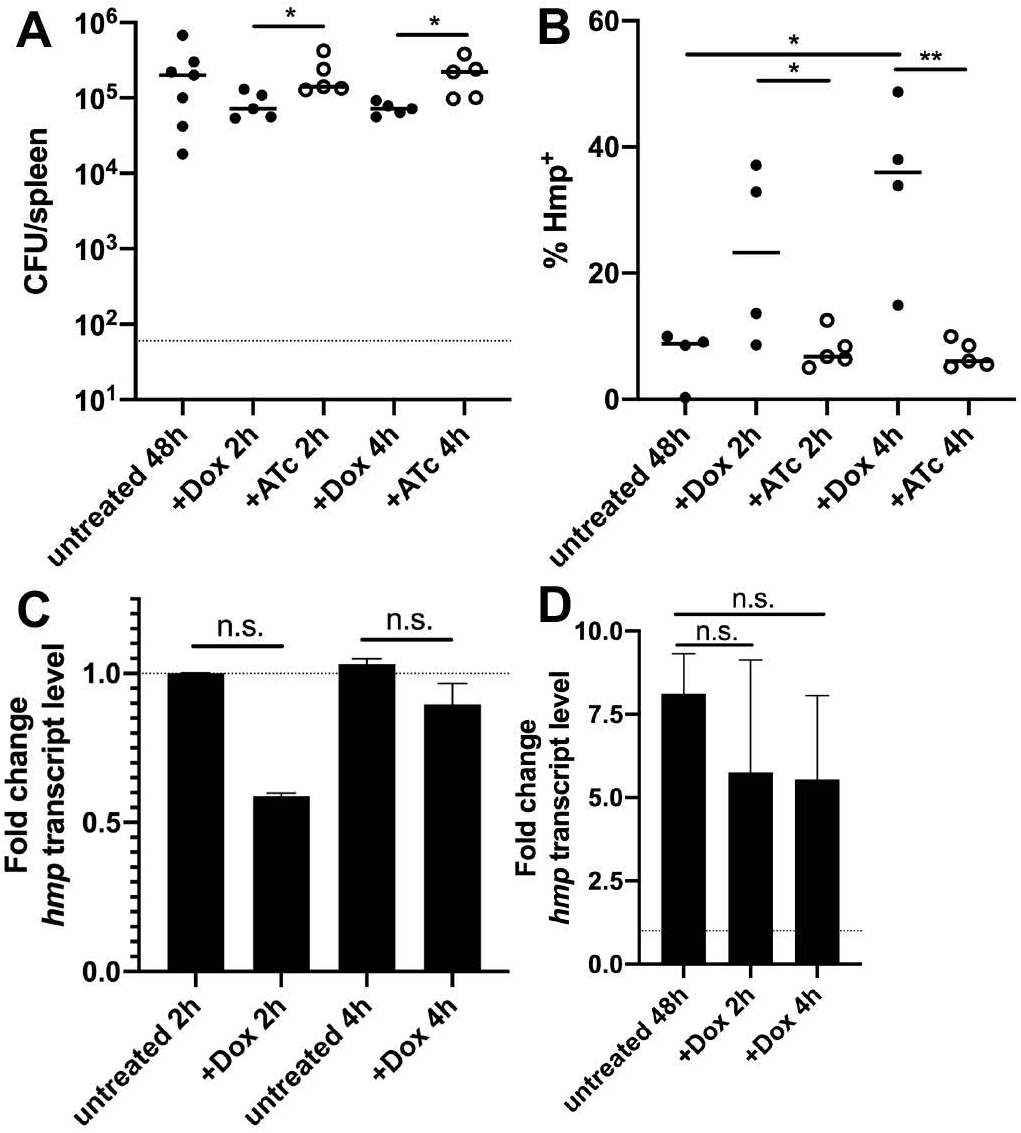
Hmp^+^ cells preferentially survive doxycycline treatment. C57BL/6 mice were inoculated i.v. with 10^3^ GFP^+^ *hmp::mCherry Y. pseudotuberculosis*. Spleens were harvested from mice at 48h p.i. (untreated). Additional mice were treated at 48h p.i with either Dox or ATc, and spleens were harvested at the indicated timepoints post-treatment to A) quantify CFUs and B) quantify the % of Hmp^+^ cells by flow cytometry. Dots: individual mice. Experiments performed in triplicate. C) WT bacteria were incubated at 37° C for 2h or 4h +/− 0.1µg/ml Dox. RNA was isolated, and qRT-PCR was used to detect *hmp* transcript levels relative to 16S. Fold change was calculated relative to untreated samples, median and range are shown for biological triplicates. D) C57BL/6 mice were infected with WT *Y. pseudotuberculosis*, and spleens were harvested from mice at 48h p.i. (untreated). Additional mice were treated at 48h p.i with Dox, spleens were harvested 2h and 4h post-treatment to isolate RNA, and perform qRT-PCR to detect *hmp* transcript levels relative to 16S. Fold change calculated relative to *hmp* levels in the inoculum. n=3 for untreated, n=4 for treated groups, median and range are shown. Statistics: A), B) & D) Kruskal-Wallis one-way ANOVA with uncorrected Dunn’s test, C) Wilcoxon matched-pairs, *p<0.05, **p<0.01. n.s.: not significant.

### Doxycycline treatment at 48h p.i. significantly prolongs mouse survival

Our results suggest that a single dose of Dox is sufficient to eliminate subsets of bacteria, but it remained unclear if bacterial numbers continue to drop after treatment, and if this single dose is ultimately sufficient to clear infection. Survival curves were performed to determine if a single dose of treatment was sufficient to prolong mouse survival and clear infection. C57BL/6 mice were infected intravenously with the WT *Y. pseudotuberculosis* GFP^+^ strain, and the infection proceeded until mice reached defined morbidity endpoints (weight loss of 15% or loss of mobility). Spleens were plated to confirm morbidity was caused by high (10^7^) CFUs. Untreated mice had a median survival of 90h (Figure 7). One Dox dose at 48h p.i. was sufficient to significantly prolong mouse survival relative to untreated mice, however, treated mice eventually succumbed to the infection with a median survival of 156h (Figure 7). Injection of Dox at 72h p.i. did not significantly prolong mouse survival (median survival: 96h), suggesting the treatment was not effective at this timepoint. These results suggest that a single dose of Dox is sufficient to significantly prolong mouse survival, but is not sufficient to promote clearance of the infection.

**Figure 7:**
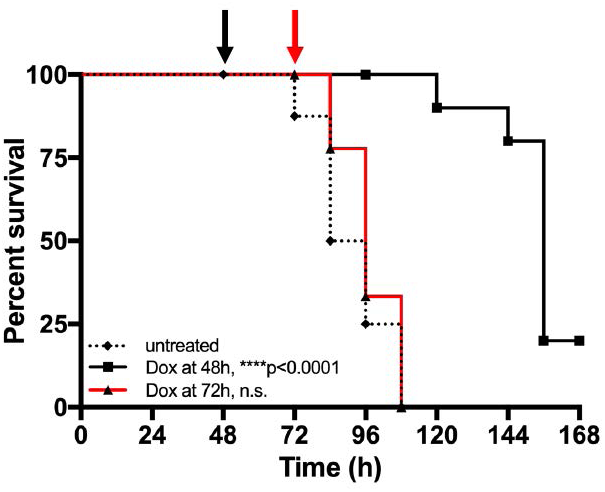
Doxycycline treatment at 48hrs significantly prolongs survival. C57BL/6 mice were inoculated i.v. with 10^3^ GFP^+^ WT *Y. pseudotuberculosis*. Mice were untreated (dotted black line) or injected with Dox at 48h p.i. (black arrow and line) or 72h p.i. (red arrow and line) and infection proceeded until mice reached morbidity endpoints. Median survival: untreated: 90h, Dox at 48h: 156h, Dox at 72h: 96h. Statistics: Log-rank (Mantel Cox) test, compared to untreated group, ****p<0.0001. n.s.: not significant.

### Bacterial growth resumes when antibiotic concentrations drop

The prolonged survival of mice in Figure 7 suggests that bacterial growth may be restricted for several days after Dox administration, but that eventually bacterial growth resumes and mice succumb to the infection. To determine when bacterial growth resumes within the spleen, C57BL/6 mice were infected with the GFP^+^ *P*_*tetA*_::*mCherry-ssrA* strain, injected with Dox at 48h p.i., and tissue was harvested at 48h and 72h post-treatment to quantify CFUs and microcolony areas (Figure 8A). CFUs remained low at 48h post-treatment, similar to 24h post-treatment (Figure 5A), however, there was a significant increase in CFUs between 48h and 72h post-treatment (Figure 8B). There was also a significant increase in microcolony areas between 48h and 72h post-treatment (Figure 8C, 8E), suggesting that bacterial growth resumes between these timepoints post-treatment.

**Figure 8:**
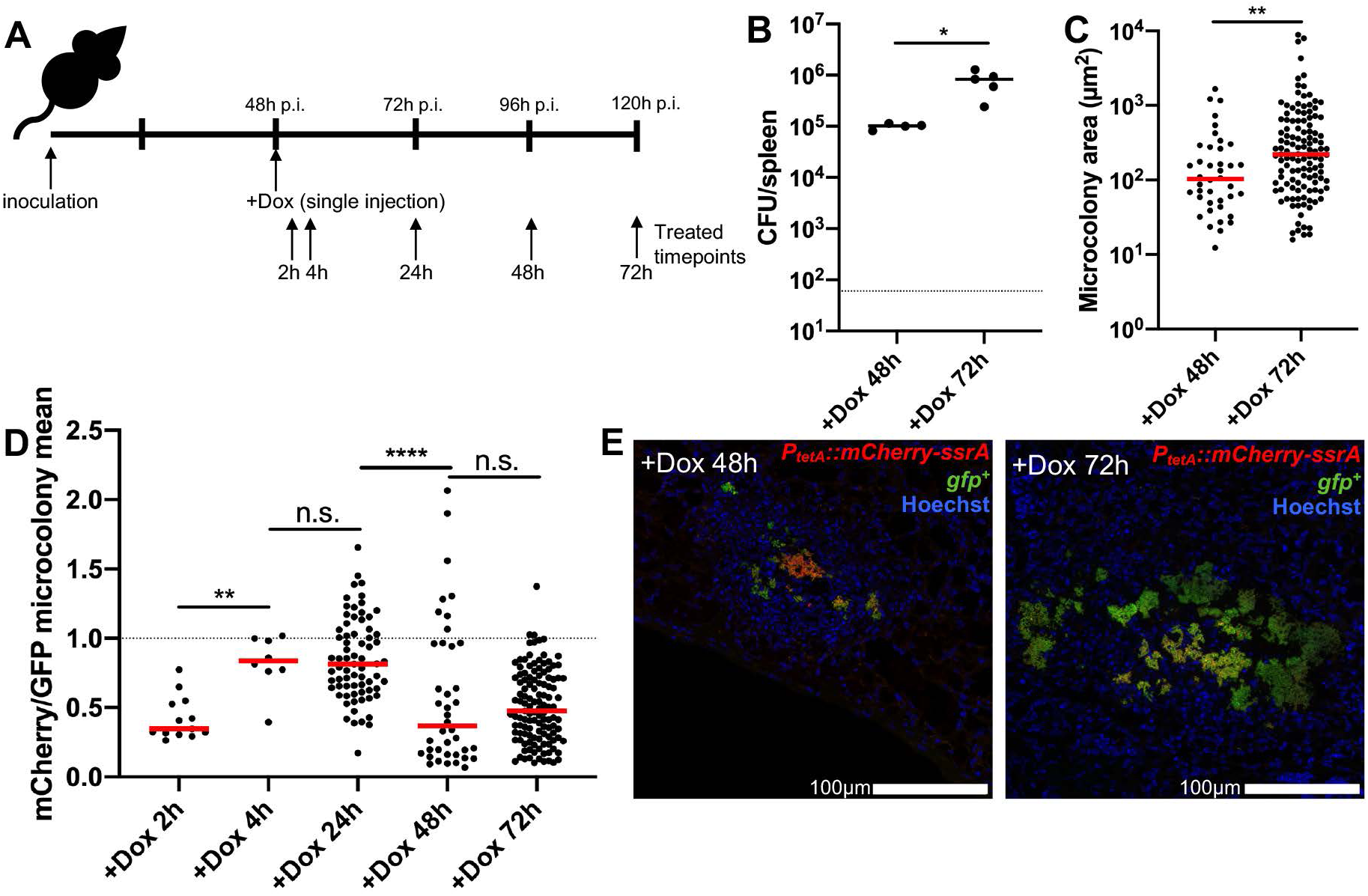
Bacterial growth resumes when antibiotic concentrations decrease. A) Timeline for mouse infections. C57BL/6 mice were inoculated i.v. with 10^3^ GFP^+^ *P*_*tetA*_::*mCherry-ssrA Y. pseudotuberculosis*. Mice were treated at 48h p.i with Dox, and spleens were harvested at the indicated timepoints post-treatment to quantify CFUs, and quantify microcolony areas and reporter expression by fluorescence microscopy. B) CFU/spleen quantification. Dots: individual mice. Represents data from two experiments. C) Microcolony area (µm^2^) quantification by fluorescence microscopy, based on GFP signal. Dots: individual microcolonies. D) Quantification of total mCherry reporter signal across microcolonies relative to total GFP signal, as a measure of antibiotic presence and exposure. E) Representative images. Medians shown. Statistics: B) & C) Mann-Whitney, D) Kruskal-Wallis one-way ANOVA with uncorrected Dunn’s test, *p<0.05, **p<0.01, ****p<0.0001. n.s.: not significant.

To determine if resumed bacterial growth correlates with a change in Dox concentration within the spleen, tissue was harvested at all indicated timepoints post-treatment to visualize Dox exposure by fluorescence microscopy (Figure 8A). Microcolonies were detected based on constitutive GFP^+^ signal, and the total mCherry signal across the microcolony was divided by the total GFP to generate a relative exposure value (mCherry mean/GFP mean fluorescence signal) for each microcolony. Using this ratio to detect exposure, there was a significant increase in antibiotic exposure between 2h and 4h post-treatment with Dox, which remained similar between 4h and 24h post-treatment (Figure 8D). Between 24h and 48h post-treatment, there was a significant decrease in Dox exposure, and this exposure value remained low at 72h post-treatment (Figure 8D). This change in Dox exposure based on reporter expression was clearly apparent within microcolonies; many microcolonies retained some Dox reporter signal at 48h post-treatment, but microcolonies were very large at 72h post-treatment and had very little remaining reporter signal (Figure 8E). These results show that Dox concentrations wane in the spleen at 48h post-treatment, and bacterial growth resumes once the concentration of Dox in the spleen has dropped to sub-inhibitory levels, ultimately causing the mice to succumb to the infection.

## Discussion

To effectively eliminate infection, antibiotics must be able to penetrate into areas of bacterial replication at high concentrations. It has been difficult to detect the concentrations of antibiotics within host tissues, and it has been assumed that treatment failure occurs because bacteria are exposed to sub-inhibitory doses, allowing bacteria to persist. Here, we show that tetracyclines diffuse into deep tissue sites and promote the elimination of a majority of the bacterial population within hours of treatment. Stressed Hmp^+^ cells preferentially survive Dox treatment, and following this initial drop in bacterial CFUs, tetracyclines accumulate at the periphery of microcolonies. This accumulation of antibiotic is sufficient to prevent bacterial growth for 48h, but then the concentration of Dox within the spleen decreases, bacterial growth resumes, and mice succumb to the infection. These results have interesting implications for other bacterial infections, and systemic infection in general, as Dox may be prescribed to treat a variety of different Gram-negative infections.

The data presented here show that Hmp^+^ cells preferentially survive Dox treatment, suggesting that stress caused by NO may be sufficient to slow bacterial growth and reduce antibiotic susceptibility. However, the Hmp^+^ peripheral population is likely exposed to multiple different host-derived stresses, and also contains a subpopulation of bacteria expressing very high levels of the T3SS (18). For this study, we used *hmp* reporter expression to mark the peripheral population, but we recognize this is likely a very complex subpopulation, and additional studies will be needed to further characterize the transcriptomic and proteomic profile of this important subpopulation. Additional studies are also needed to truly show Hmp^+^ cells represent a slow-growing subpopulation with decreased metabolic activity, however the decreased antibiotic susceptibility of this population suggests this is likely the case (5, 6, 8).

During Dox treatment, many bacterial cells survive and are capable of resuming growth once antibiotic concentrations wane (Figure 8). Our data suggests that bacteria may be exposed to Dox concentrations ranging from 0.1-1µg/ml for as long as 48h, and it will be very interesting to determine how this prolonged exposure impacts the bacterial population. The genes expressed in response to Dox could be potential targets to more effectively eliminate the bacteria population with combination therapeutics. We would expect that many of the genes expressed during this long term Dox exposure are protective responses, that if inhibited, would result in a loss of bacterial viability.

Doxycycline treatment has direct effects on the bacterial population, but the release of bacterial products at the site of infection will also impact the host response. Our experiments to detect preferential survival of the Hmp^+^ population were focused on early timepoints post-treatment, in part because we were interested in early perturbation by Dox, but also because interactions between bacteria and host cells are likely very different at later hours post-treatment as the host immune response becomes activated by release of bacterial ligands. As additional, activated immune cells subsets are recruited to the spleen, we expect the bacterial population to respond to their presence. It would be very interesting to further characterize the bacterial and host response at later timepoints after Dox treatment, which may provide important clues as to why the host response was not sufficient to completely eliminate the bacterial population.

Antibiotic susceptibility is typically assessed with *in vitro* experiments, and very few studies have focused on the impact of antibiotics in the host environment, where bacteria may respond very differently, and may have very different susceptibility patterns. It is critical that we develop tools, such as the fluorescent reporter developed here, to study antibiotic activity, diffusion, and efficacy within mammalian tissues.

## Materials and Methods

### Bacterial strains & growth conditions

The WT *Y. pseudotuberculosis* strain, IP2666, was used throughout (18, 33). For all mouse infection experiments, bacteria were grown overnight (16h) to post-exponential phase in 2xYT broth (LB, with 2x yeast extract and tryptone) at 26°C with rotation. For antibiotic exposure experiments, bacteria were grown in LB overnight (16h) at 26°C with rotation, sub-cultured 1:100, and grown at 37°C with rotation in the absence or presence of the indicated concentrations of doxycycline (Dox) or anhydrotetracycline (ATc), for the indicated timepoints.

### Murine model of systemic infection

Six to 8-week old female C57BL/6 mice were obtained from Jackson Laboratories (Bar Harbor, ME). All animal studies were approved by the Institutional Animal Care and Use Committee of Johns Hopkins University. Mice were injected intravenously into the tail vein with 10^3^ bacteria for all experiments. At the indicated timepoints post-inoculation (p.i.), spleens were removed and processed. Intact tissue was stabilized for RNA isolation or fixed for fluorescence microscopy. Tissue was homogenized to quantify CFUs and process for flow cytometry. For survival curves, the following morbidity endpoints were used: weight loss of 15% or greater (measured initially and every 24 hours p.i.), lethargy, hunched stature, or reduced activity; any of these endpoints qualified mice for euthanasia. Spleens were plated to confirm morbidity was associated with high bacterial CFUs.

### Generation of reporter strains

Two of the *Y. pseudotuberculosis* reporter strains in this study were previously described: WT GFP^+^ and WT GFP^+^ *hmp::mCherry* (18). GFP^+^ strains were constructed by transformation with the constitutive GFP plasmid, which expresses GFP from the unrepressed *Ptet* of pACYC184 (18, 20). The destabilized version of *mCherry* was constructed by fusing an 11 amino acid *ssrA* tag, with AAV terminal amino acids, between the last coding amino acid of *mCherry* and the stop codon (29, 30). This was inserted downstream of the *P*_*tac*_ IPTG inducible promoter in pMMB207∆267 (34) to quantify mCherry half-life following IPTG induction. A stable, WT *mCherry* construct was generated in parallel as a control. The *P*_*tetA*_::*mCherry* reporter was constructed by amplifying *tetR* through *P*_*tetA*_ promoter sequence from Tn10, fusing this to *mCherry* by overlap extension PCR to replace *tetABC* with *mCherry*, and cloning this into the low copy pMMB67EH plasmid. *P*_*tetA*_::*mCherry*-*ssrA* was constructed the same way, by amplifying the *ssrA*-tagged, destabilized *mCherry*.

### Fluorescence detection, plate reader

Bacteria were washed in PBS, resuspended in equivalent volume PBS, then added to black walled, clear bottom 96 well plates. Optical density (A_600nm_) was used to approximate cell number; mCherry fluorescence was detected with 560nm excitation/610nm emission, using a Synergy H1 microplate reader (Biotek).

### Determining fluorescent protein half-life

Overnight cultures (16h) were grown in LB with 1mM IPTG. Bacteria were washed 3x in PBS to remove IPTG, resuspended in PBS, and kanamycin (50µg/ml) was added to inhibit additional protein translation. Bacteria were added to black walled, clear bottom 96 well plates, and were incubated in a Tecan Infinite M200 (Tecan) microplate reader for the indicated timepoints. Fluorescence reads (580nm excitation, 620nm emission) were taken every 5 minutes. 560nm excitation/610nm emission detection with a Synergy H1 microplate reader (Biotek) gave equivalent results. Fluorescent protein half-life was calculated from six biological replicates using a one-phase exponential decay equation in Prism software.

### Fluorescence microscopy: bacteria

To visualize individual bacterial cells, samples were pelleted, resuspended in 4% paraformaldehyde (PFA) and incubated overnight at 4° C for fixation. PFA was removed and bacteria were resuspended in PBS prior to imaging. Agarose pads were prepared to immobilize bacteria for imaging, by solidifying a thin layer of 25µl 1% agarose in PBS between a microscope slide and coverslip. Once solidified, coverslips were removed, bacteria were added, coverslips were replaced, and bacteria were imaged with the 63x oil immersion objective, using a Zeiss Axio Observer 7 (Zeiss) inverted fluorescent microscope with XCite 120 LED boost system and an Axiocam 702 mono camera (Zeiss). Volocity image analysis software was used to specifically select individual bacterial cells and quantify the fluorescent signal associated with each cell.

### Fluorescence microscopy: host tissues

Spleens were harvested and immediately fixed in 4% PFA in PBS for 3 hours. Tissues were frozen-embedded in Sub Xero freezing media (Mercedes Medical) and cut by cryostat microtome into 10µm sections. To visualize reporters, sections were thawed in PBS, stained with Hoechst at a 1:10,000 dilution, washed in PBS, and coverslips were mounted using ProLong Gold (Life Technologies). Tissue was imaged as described above, with an Apotome.2 (Zeiss) for optical sectioning.

### qRT-PCR to detect bacterial transcripts in broth-grown cultures

Bacterial cells were grown in the presence or absence of 0.1µg/ml Dox for the indicated time points, pelleted, resuspended in Buffer RLT (QIAGEN) + ß-mercaptoethanol, and RNA was isolated using the RNeasy kit (QIAGEN). DNA contamination was eliminated using the DNA-free kit (Ambion). RNA was reverse transcribed using M-MLV reverse transcriptase (Invitrogen), in the presence of the RNase inhibitor, RnaseOUT (Invitrogen). Approximately 30 ng cDNA was used as a template in reactions with 0.5 µM of forward and reverse primers (18) and SYBR Green (Applied Biosystems). Control samples were prepared that lacked M-MLV, to confirm DNA was eliminated from samples. Reactions were carried out using the StepOnePlus Real-Time PCR system, and relative comparisons were obtained using the ∆∆CT or 2^−∆Ct^ method (Applied Biosystems). Kits were used according to manufacturers’ protocols.

### qRT-PCR to detect bacterial transcripts from mouse tissues

Spleens were harvested, and immediately submerged in RNALater solution (QIAGEN). Tissue was homogenized in Buffer RLT + ß-mercaptoethanol, and RNA was isolated using the RNeasy kit (QIAGEN). Bacterial RNA was enriched following depletion of host mRNA and rRNA from total RNA samples, using the MICROB*Enrich* kit (Ambion). Kits were used according to manufacturers’ protocols; DNA digestion, reverse transcription, and qRT-PCR were performed as described above.

### Flow cytometry

Homogenized tissue was treated with collagenase D and DNase I in PBS + 0.2mM EDTA to dissociate host cells from bacteria. Host cells were lysed with eukaryotic lysis buffer (5mM EDTA, 0.1% NP-40, 0.15M NaCl, 1M HEPES), debris settled out of solution, supernatant was collected, pelleted and fixed in 4% PFA overnight at 4° C. Bacteria were detected using GFP signal and FSC (size), and 1000 bacterial events were collected per tissue, using a BD FACSCalibur. For analysis, bacterial cells were gated based on GFP signal, and GFP^+^ bacterial controls lacking mCherry constructs (prepared and collected in parallel) were used to identify mCherry^+^ cells.

### Image analysis

Volocity image analysis software was used to quantify microcolony areas and fluorescence as previously described (18). Zen2 co-localization software was used to generate an intensity dot plot for pixels within an image (Figure 2F) with a corresponding pseudocolor image; each quadrant of the dot plot corresponds with a highlighted pseudocolor. Bacterial cells defined the GFP^+^ threshold; within this threshold, the pixels with the highest mCherry signal intensity (66%) were considered mCherry^high^, the lowest (34%) mCherry signal intensity was considered mCherry^low^. Image J was used to quantify the signal intensity of each channel at the centroid and periphery of each microcolony, to generate relative signal intensities of fluorescent reporters, as previously described (18). Thresholding was used to define the total area and centroid of each microcolony, and 0.01 pixel^2^ squares were selected to calculate values. Peripheral measurements depict bacteria in contact with host cells; peripheral cells are highlighted outside dotted lines in Figures 2E, 3E & 3F.

## Author Contributions

Designed and performed experiments: JRR, ALE, KMD. Intellectual/conceptual contribution: JRR, KMD. Analyzed the data: JRR, KMD. Wrote the manuscript: ALE, BL, KMD.

## Acknowledgments

We thank the members of the Davis lab for constructive feedback and suggestions during manuscript preparation. We thank Ralph Isberg for invaluable feedback throughout. The authors of this manuscript declare no conflicts of interest. This work was supported by a NIAID K22 Career Transition Award (1K22AI123465-01).

---

**Supplemental Figure 1:**
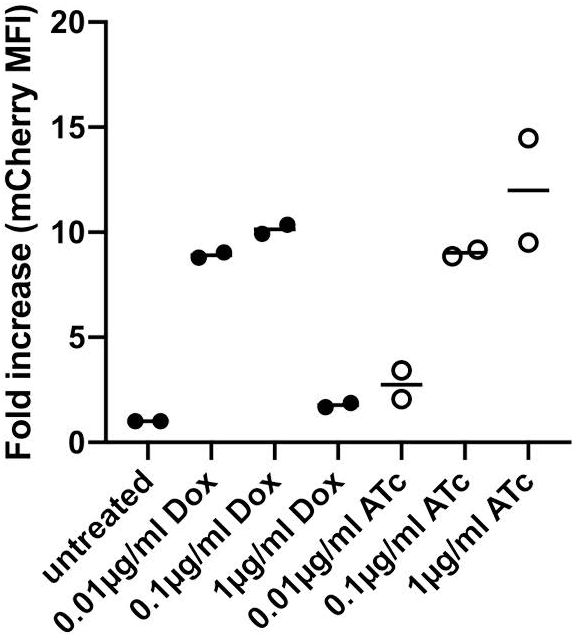
Doxycycline-inhibited cells have low levels of mCherry signal. The *P*_*tetA*_::*mCherry* strain was exposed to the indicated doses of doxycycline (Dox) or anhydrotetracycline (ATc) during growth in LB at 37°C. mCherry reporter expression within individual bacterial cells was detected by flow cytometry after 4h treatment, and is expressed as the fold increase in mCherry mean fluorescent intensity (MFI) relative to the signal from untreated cells. Experiment performed in duplicate, biological replicates shown with medians.

